# Identification of pathways modulating Vemurafenib resistance in melanoma cells via a genome-wide CRISPR/Cas9 screen

**DOI:** 10.1101/2020.07.11.198911

**Authors:** Corinna Jie Hui Goh, Jin Huei Wong, Chadi El Farran, Ban Xiong Tan, Cynthia Coffill, Yuin-Hain Loh, David Lane, Prakash Arumugam

## Abstract

Vemurafenib is a BRAF kinase inhibitor (BRAFi) that is used to treat melanoma patients harbouring the constitutively active BRAF-V600E mutation. However, after a few months of treatment patients often develop resistance to vemurafenib leading to disease progression. Sequence analysis of drug-resistant tumour cells and functional genomic screens have identified several genes that regulate vemurafenib resistance. Reactivation of mitogen-activated protein kinase (MAPK) pathway is a recurrent feature of cells that develop resistance to vemurafenib. We performed a genome-scale CRISPR-based knockout screen to identify modulators of vemurafenib resistance in melanoma cells with a highly improved CRISPR sgRNA library called Brunello. We identified 33 genes that regulate resistance to vemurafenib out of which 14 genes have not been reported before. Gene Ontology enrichment analysis showed that the hit genes regulate histone modification, transcription and cell cycle. We discuss how inactivation of hit genes might confer resistance to vemurafenib and provide a framework for follow-up investigations.

## INTRODUCTION

Vemurafenib, also known as PLX4032, is one of the first FDA (Food and Drug Administration)-approved small molecule inhibitors for the treatment of metastatic melanoma patients, specifically for patients harbouring the BRAF-V600E mutation. The BRAF-V600E mutation is present in approximately 50% of melanoma cases and causes constitutive activation of BRAF and the MAPK (ERK) signalling pathway leading to uncontrolled cell proliferation [1]. Vemurafenib binds specifically to the adenosine triphosphate (ATP) binding pocket of activated BRAF-V600E, blocks ERK1/2 activation, and induces cell cycle arrest and apoptosis [2]. Although there is short-term tumour regression and enhancement of patient survival, resistance to vemurafenib frequently develops [1]. Therefore, an understanding of mechanisms underlying vemurafenib resistance is of paramount importance to develop novel melanoma therapeutics.

Exposure of melanoma cell lines to incremental concentrations of vemurafenib resulted in drug resistance [3]. In contrast, intermittent use of vemurafenib was observed to delay acquisition of resistance. Possibly, the continuous administration of vemurafenib provides a selective pressure for drug-resistant cells to thrive [4]. Initial inhibition of BRAF-V600E by vemurafenib severely depletes MAPK output and is therefore considered to be a dormant period for resistant cells to accumulate before resulting in their over proliferation. On the other hand, excessive MAPK output causes toxicity [4], [5]. Hence, the need to re-establish dynamic equilibrium with interrupted scheduled dosing of vemurafenib was emphasized [6].

Basal phosphorylation levels of both MEK and ERK were enhanced in vemurafenib-resistant A375 cells while AKT phosphorylation levels remained relatively unchanged [7]. Phosphorylation of downstream effectors of mTOR (decreased phospho-S6 and hyperphosphorylation of 4E-BP1) has also been shown to contribute to vemurafenib resistance [8]. Reactivation of MAPK pathway and/or activation of PI3K/AKT pathways confer resistance to BRAFi in melanoma cells [9]. MAPK activation could occur by increased expression of their upstream activators namely the Receptor Tyrosine Kinases. Alternatively, activating mutations within the RAS/RAF/MEK/Erk signalling pathway could recover the MAPK pathway [9]. Sequence analysis of vemurafenib-resistant cancer cells have identified mutations in BRAF-V600E (amplification, truncation, alternative splicing and fusions), activating mutations in RAS genes (NRAS, KRAS, HRAS), RAC1, MAP2K1 and AKT and loss-of-function mutations in GNAQ/GNA11, CDKN2A, PTEN, PIK3R2 and DUSP4. However, the molecular basis of about 30-40% cases of BRAFi resistance remains unknown.

Functional genetic screens have been gaining traction as a powerful approach to identify novel cellular pathways involved in acquisition of drug-resistance. Whittaker *et al.* (2013) used genome-scale RNA interference screen with a library size of 90,000 shRNAs targeting approximately 16,600 genes expressed in A375 cells [10]. Seminal work on the genome-wide CRISPR KnockOut (GeCKO) screen [11] for regulators for vemurafenib resistance was performed with two half sgRNA GeCKOv1 libraries, covering 18,000 human genes with a total of 6 target sgRNAs per gene. Doench *et al.* (2016) and Sage *et al.* (2017) used GeCKOv2 libraries to screen for the vemurafenib resistant A375 cells [12], [13]. Sanson *et al.* (2018) used Calabrese and SAM libraries to screen for the vemurafenib resistant A375 cells [14]. Although some genes like NF1, TAF6L and CUL3 were obtained in all the screens, the extent of overlap between the different genetic screens was limited.

A second-generation sgRNA library called ‘Brunello’ was designed by optimizing targeting efficiency and minimizing off-target effects. Brunello outperformed the first-generation GeCKO library [14]. In this study, we performed a CRISPR-Cas9 mediated genome-wide knockout screen in human melanoma cell line A375 using the Brunello library to identify novel genes that regulate vemurafenib resistance. Gene ontology enrichment analysis of the top hits suggest that alterations to MAPK pathway, epigenome and cell cycle facilitate acquisition of resistance to vemurafenib in melanoma cells.

## MATERIALS AND METHODS

### Library amplification

100 ng of human sgRNA library Brunello in lentiCRISPRv2 (Addgene, #73179) was transformed into electrocompetent Endura cells (#60242, Lucigen) in quadruplicates. Cells were transferred into a chilled electroporation cuvette and pulsed (BioRad Micropulser, #165-1200) at 10 μF, 600 Ω, 1800 V. Within 10 s of the pulse, 1975 μL of the Recovery medium was added to the cells. Cells were then incubated in a shaker incubator at 250 rpm for 1 h at 37°C before being selected on LB-Agar containing 100 μg/ml ampicillin in 245 mm Square BioAssay Dishes (Corning) at 32°C for 14 h. Transformants were pooled by rinsing the bioassay plates with 20 mL of LB twice using a cell scraper and used for plasmid DNA extraction with Endotoxin-free plasmid DNA purification (#*740424-10*, Macherey Nagel).

### Lentivirus generation and harvesting

Lipofectamine transfection was performed on HEK293FT cells 24 h after the cells were seeded into 100 mm dishes in DMEM (Dulbecco’s Modified Eagle’s Medium media) supplemented with 10% Fetal Bovine Serum (FBS) (#SH30071.03, Hyclone), 2 mM L-glutamine and 1 mM sodium pyruvate. Briefly, 612.5 μL of Opti-MEM containing P3000 (14 μL), DNA mixture of lentivirus packing plasmid (2 μg PLP1, 2 μg PLP2, 1 μg pVSVG) and Brunello library (2 μg) were added into Lipofectamine mixture (612.5 μL of Opti-MEM and 14 μL of Lipofectamine 3000). This mixture was incubated at room temperature for 15 min. Afterwards, the transfection mixture was added dropwise to the HEK293FT cells and incubated in the 37 °C incubator with 5% CO2 (w/v). After 24 h, media was supplemented with 1 mM sodium butyrate to increase virus production. For the next two consecutive days, the viruses were harvested by centrifugation at 16,500 rpm for 90 minutes at 4°C. Virus pellets were re-suspended in PBS and kept at −80°C. To determine multiplicity of infection (MOI) of Brunello virus, a serial dilution of the virus at 1:10, 1:50, 1:100, 1:1000 was added into 1 × 10^6^ A375 cells in a 12-well plate. Polybrene was then added at a final concentration of 5 μg/mL and then centrifuged at 3,000 rpm for 90 minutes at 30°C. After 1 day of incubation, we equally split each virus transduced cells into two sets of medium with or without 1 μg/ml puromycin. After 3 days of incubation, cell numbers (*n*) were determined per condition and the percentage transduction was calculated as followed:

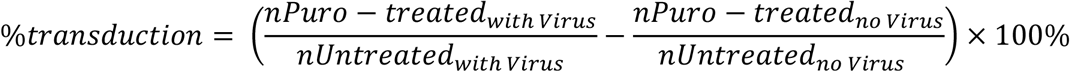

A MOI of 0.4 was used for large-scale screening.

### Determination of IC50 for Vemurafenib in A375

IC50 was determined on days 3 and 4 after vemurafenib treatment using Real-time-Glo MT Cell Viability Assay according to the manufacturer’s protocol (Promega, #G9711). Using a 96-well white plate, viability assays of 2000 A375 cells were performed in triplicates. IC50 for vemurafenib was determined with a dose-response curve.

### Vemurafenib resistance screen with the Brunello library

9.6 × 10^7^ A375 cells were transduced with 1 × 10^6^ cells plated per transduction well (12-well plate). 110 μL of the concentrated Brunello library (MOI=0.4) was applied into each well containing A375 cells. Puromycin (1 μg/mL) was added to the cells 24 hours post transduction and maintained for 7 days. On Day 7, cells were split into DMSO or 2μM vemurafenib conditions in duplicates with 3 × 10^7^ cells per replicate and an additional 3.3 × 10^7^ cells were frozen down for Brunello library genomic DNA analysis. DMSO-treated cells were passaged every 3-4 days. Fresh medium with 2 μM vemurafenib was added to drug-treated cells every 3-4 days. Cell pellets were taken at 7 days and 14 days after vemurafenib treatment.

### Genomic DNA (gDNA) extraction & Amplicon sequencing

Cell pellets were thawed for genomic DNA extraction using Blood & Cell Culture DNA Midi/Maxi Kit (Qiagen). PCR was performed with Q5 Hot Start High-Fidelity 2 × Master Mix (#M0494L, New England Biolabs) with the amount of input genomic DNA (gDNA) for each samples was 15 μg. For each sample, we performed 10 separate 50 μL reactions with 1.5 μg genomic DNA, in each reaction using 10 forward primers with 1-10 bp staggered region to increase the diversity of the library and 1 reverse primer containing the unique barcode to differentiate the samples (Table S1). Thermal cycling conditions were set up with the following:

| Cycle# | Denaturation | Annealing | Extension |
| --- | --- | --- | --- |
| 1 | 98°C, 3 min |  |  |
| 2-29 | 98°C, 10 s | 60°C, 30 s | 72°C, 25 s |
| 30 |  |  | 72°C, 5 min |

Minimum number of PCR cycles to amplify the gDNA was determined by performing small-scale PCRs with varying number of cycles. All 10 PCRs of a single sample were pooled and purified using QIAquick PCR Purification Kit (Qiagen). The purified PCR product was then separated using 2% agarose gel and gel extracted with the Qiagen Gel Extraction Kit. These gel purified PCR products were stored at −20°C before being submitted for next-generation sequencing (NGS) by NovogeneAIT Genomics (Singapore) using HiSeq-SE150 platform.

### Data analyses

The raw FASTQ files (.fastq.gz) were demultiplexed by NovogeneAIT Genomics (Singapore) and then uploaded to CRISPRAnalyzer (http://www.crispr-analyzer.org) [15] for analysis of phenotypes against the reference Brunello library. CRISPRAnalyzer performs the alignment of the unique sgRNA sequence with the reference library to obtain the sgRNA read counts. After extraction of read count files (.txt), these files were uploaded again to CRISPRAnalyzer. To determine the log_2_ fold changes of gRNA read counts between treatment and control conditions, read counts with less than 20 sgRNAs were removed for analysis. Model-based Analysis of Genome-wide CRISPR/Cas9 Knockout (MAGeCK) was chosen for our analysis which scores whether each gene is enriched or depleted and generates a gene ranking list [16]. Default settings were used in our analysis. Genes with P values less than 0.05 were chosen for further analysis [17]. Shortlisted genes were analyzed using the Search Tool for the Retrieval of Interacting Genes/Proteins database (STRING) [18] for functional association. The text mining option was disabled during PPI data analysis to ensure higher confidence. Gene Ontology (GO) term enrichment analysis was performed using DAVID [19][20].

## RESULTS AND DISCUSSION

### Functional interrogation of genes in modulation of Vemurafenib resistance via CRISPR

We sought to identify novel regulators of vemurafenib resistance using the highly optimized second-generation Brunello CRISPR sgRNA library. We chose A375 melanoma cells which harbour the gain-of-function BRAF-V600E mutation. Treatment of A375 cells with vemurafenib resulted in a growth arrest with an IC50 of 248.3 nM (Figure 2A).

To select for vemurafenib-resistant A375 cells in our genome wide CRISPR screen, we used vemurafenib at 2 μM, which is about 10-fold higher than its IC50 value. We transduced A375 cells with the Brunello lentiviral sgRNA library at a MOI of 0.4 and selected for transductants in the presence of puromycin for 7 days. We then harvested the puromycin-resistant cells and treated them with either DMSO or 2 μM vemurafenib for 7 days and 14 days in duplicates (Figure 1). We isolated the genomic DNA from cells collected at Day 0, Day 7 (DMSO or vemurafenib-treated) and Day 14 (DMSO or vemurafenib-treated), amplified the sgRNA sequences from genomic DNA by PCR and sequenced the PCR products by deep sequencing methods.

**Figure 1:**
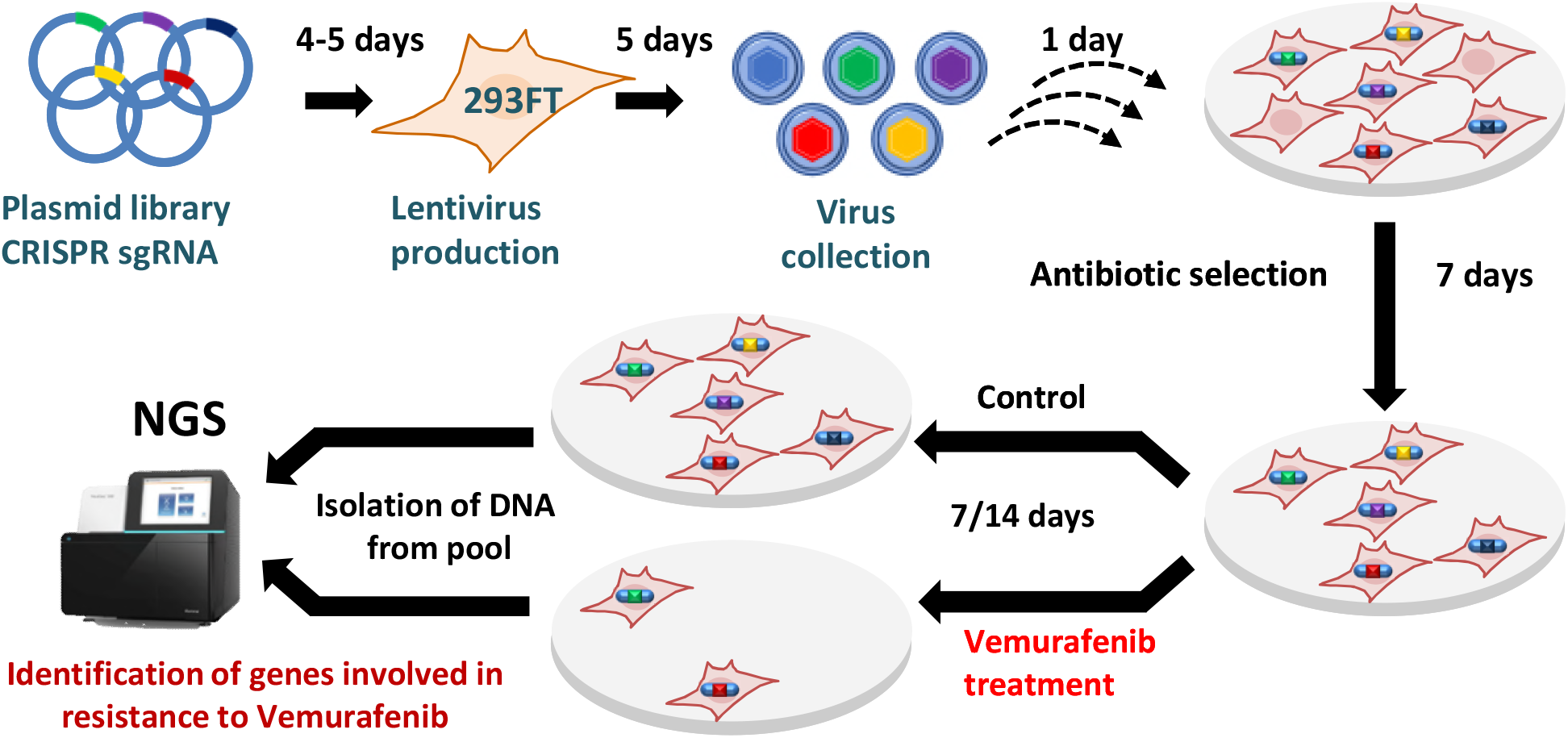
Schematic representation of the workflow for the genome-wide CRISPR/Cas9 screen. HEK293FT cells were transfected with lentiviral CRISPR-Cas9 plasmid library (Brunello library with genome wide coverage and 4 sgRNAs per gene) for lentivirus production. A375 cells were then transduced with lentiviruses generated from the Brunello plasmid library. Cells were selected for successful transduction using puromycin selection for 7 days. After that, cells were either treated with 2 μM vemurafenib or DMSO. Samples were collected at days 7 and Genomic DNA was prepared from the cells and the relative sgRNA abundance was determined by PCR-mediated amplification of sgRNA sequences from genomic DNA followed by NGS.

CRISPRAnalyzer–based analysis of NGS data showed that less than 7.5% of the Brunello library sgRNAs were covered by less than 100 reads (Figure 2B, Table S2). This meant that 92.5% of the Brunello library sgRNA have a read count of more than 100, indicating a good library coverage. In terms of sgRNA distribution, Day 14 demonstrated a higher difference between vemurafenib-treated cell and control in comparison with Day 7 (Figure 2C). The sgRNA distribution in Day 14 vemurafenib-treated cells (P1D14 and P2D14) was significantly different from sgRNA distribution in DMSO-treated cells (C1D14 and C2D14) (Figure 2C). Day 14 vemurafenib-treated cells showed a higher range of read counts indicating an enrichment of some sgRNA-treated cells after drug treatment (Figure 2C). Analysis of the pairwise correlation data indicated that the replicates of the Day 7 treatment correlated well with a Pearson value of 1 and Spearman value of 0.93 indicating good reproducibility (Figure 2D). For the day 14 replicates, DMSO-treated samples (C1D14 and C2D14) correlated well (Pearson value = 1, Spearman = 0.95) but for the vemurafenib-treated samples (P1D14 and P2D14), their correlation was relatively low (Pearson value = 0.68, Spearman = 0.82) (Figure 2E). This suggests that our genetic screen was not saturating and some modulators of vemurafenib resistance may have been missed out. However, the altered read count distribution of vemurafenib–resistant cells in comparison to DMSO-treated cells indicated that our screen has identified mutations that enhance resistance to vemurafenib.

**Figure 2:**
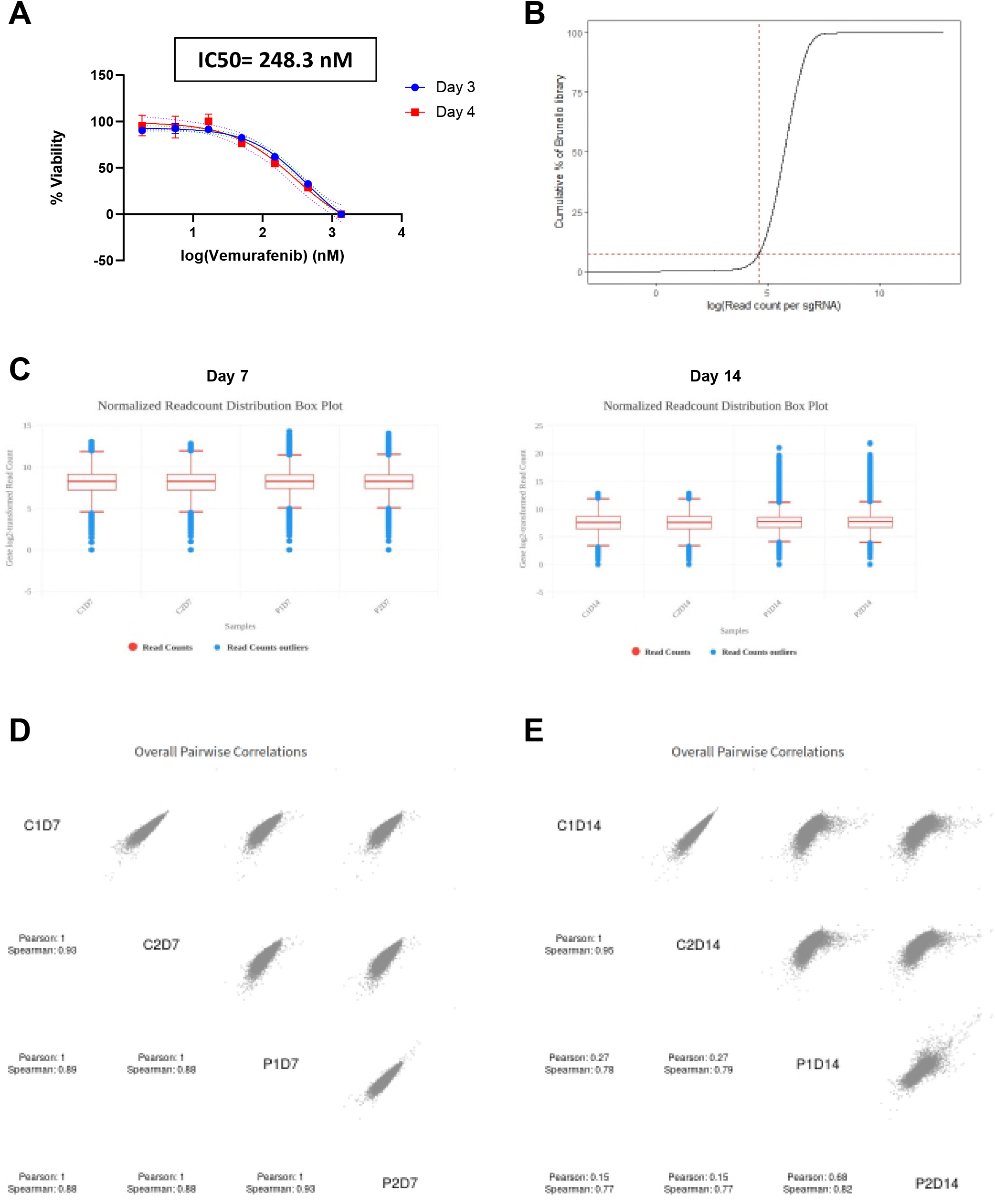
NGS data analysis from the genome wide screen using CRISPRAnalyzeR. A. Cell viability of A375 cells after treatment with various concentration of vemurafenib was determined using Real-time-Glo MT Cell Viability Assay. % Viability was plotted against various concentration of vemurafenib. 3 days after vemurafenib treatment samples are indicated by circle shape while the 4 days after vemurafenib treatment samples are indicated by square shape. B. Cumulative distribution of the number of reads per sgRNA in a single A375 experiment. The dotted line indicates that less than 7.5% of the sgRNAs are covered by less than 100 reads. C. Boxplot showing the read count distribution from individual sgRNAs for the DMSO-treated and vemurafenib-treated cells for day 7 and day 14. For day 14, there is an increase in the number of reads for the most abundant sgRNAs in the vemurafenib-treated cells as compared to the DMSO treated cells. D. Pairwise correlation plot of the Day 7 samples with the Pearson and Spearman correlation coefficients. E. Pairwise correlation plot of the Day 14 samples with the Pearson and Spearman correlation coefficients.

### Vemurafenib resistance genes

Using the CRISPRanalyzer platform to assess our NGS data, we identified 33 genes (p-value <0.05) that had significantly enriched sgRNA levels after 14 days of vemurafenib treatment (Figure 3 and Table S3). All the top 10 hits have been reported in previous genome-wide CRISPR/Cas9 screens validating our screen for modulators of vemurafenib resistance. The top 10 enriched genes were CCDC101 (SAGA complex associated factor 29), TAF6L (TATA-box binding protein associated factor 6 like), SUPT20H (Transcription factor SPT20 homolog, SAGA complex component), TADA2B (Transcriptional Adaptor 2B), NF2 (Neurofibromin 2), MED12 (Mediator complex subunit 12), TADA3 (Transcriptional adaptor 3), CUL3 (Cullin 3), TADA1 (Transcriptional adaptor 1) and MED23 (Mediator Complex Subunit 23).

**Figure 3:**
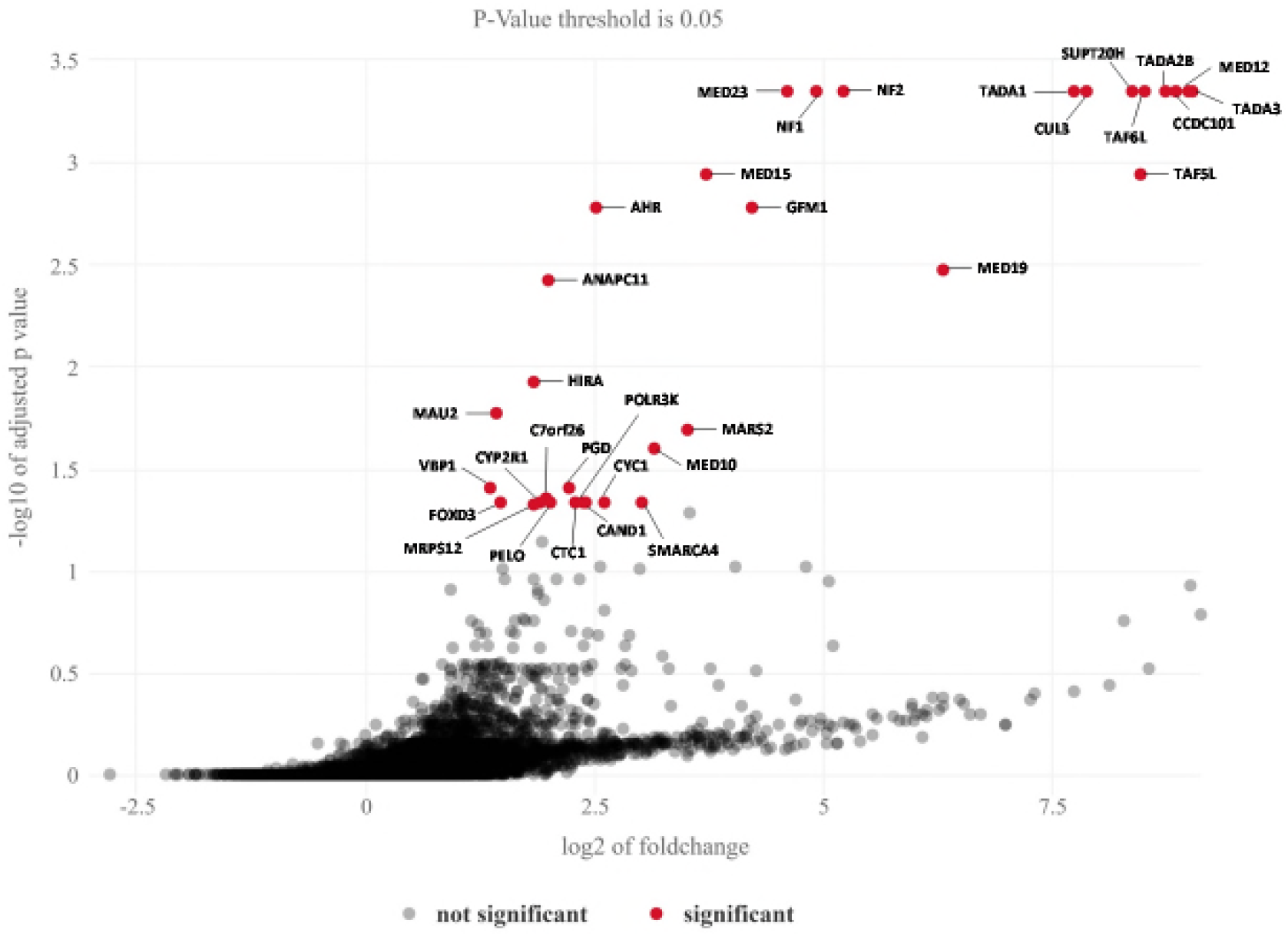
Identification of vemurafenib-resistance genes using MAGeCK. The sgRNA distribution data of DMSO-treated and vemurafenib-treated cells were analyzed by MaGeCK. For each gene, the −log_10_ P-value was plotted against its log_2_ fold change. Vemurafenib resistance genes were identified using a P-value threshold of 0.05. Significant hits are denoted by red dots along with gene names.

### Comparison with previous genetic screens

Four genome wide screens (1 shRNA, 2 CRISPRko and 1 CRISPRi) for regulators of vemurafenib resistance have been reported previously [12][13][16][11][10]. We compared our top 33 hits (p value <0.05) with Top 100 enriched hits obtained from four previous genome wide screens [12][13][16][11][10]. Out of our top 33 hits, only 19 were obtained in previous studies (Table S4). We also compared our CRISPRko Brunello library top 33 enriched hits with top 100 enriched hits from the two previous CRISPRko screens. As depicted in the Venn diagram (Figure S1), only 13 hits (NF1, NF2, MED12, MED15, MED19, MED23, CUL3, TADA1, TADA2B, CCDC101, TAF5L, TAF6L, PGD) were obtained in all three screens. These results indicate that there is considerable variability with results obtained from independent CRISPR screens. Fourteen out of the 33 hits have not been previously reported in genome wide screens.

To assess the presence of functional associations between proteins that confer vemurafenib resistance, we submitted our top 33 hits to the STRING v11database [18] to construct the Protein-Protein interaction (PPI) network. Proteins in the PPI network do not need to physically interact but they may sufficiently overlap in their functional roles and participate in a common biological process [18]. PPI network among the hits is shown in Figure 4A, which includes a total of 33 nodes and 38 edges, with the node and edge representing a target protein and PPI respectively. The average node degree is 2.3 which represents the average number of targets connected to a target. Greater the degree of a target, the stronger is its role in the interaction network. Our hits interact with each other significantly with a high confidence score (average combined associated score of 0.96 and PPI enrichment p-value: < 1.0e-16) (Table S5). MED12 interacts with MED10, MED15, MED19 and MED23 with a combined associated score of 0.999 (Figure 4A and Table S5). The combined associated score of TADA1 with SUPT20H, TADA2B, TADA3, CDC101, TAF5L and TAF6L lies between 0.981 and 0. 997 (Figure 4A and Table S5). Presence of a strong functional association between the hits demonstrates the robustness of the results from the genome-scale screen.

**Figure 4:**
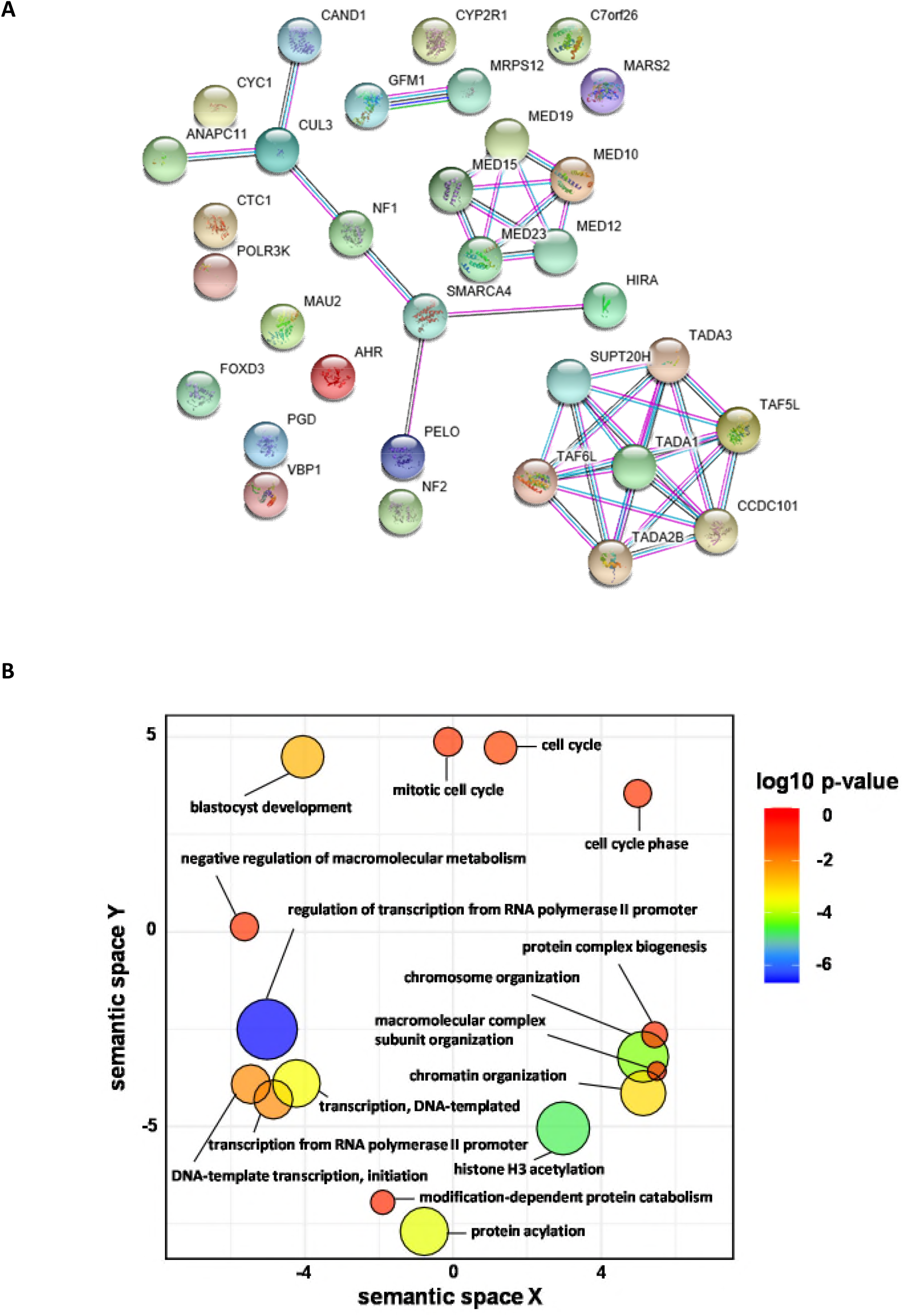
STRING analysis and Gene ontology term enrichment analysis of the 33 hits from the CRISPR screen. A. Shortlisted genes were analyzed using the Search Tool for the Retrieval of Interacting Genes/Proteins database (STRING v11). Network nodes represent proteins and the edges represent protein-protein interactions. Results of STRING analysis are presented in Table S5. B. GO term enrichment analysis was performed using DAVID. GO terms are clustered together based on semantic similarity in the 2-dimensional scatterplot. P-value which indicates the enrichment strength in the annotation category is denoted by the bubble colour and the GO term frequency is denoted by the bubble size.

To gain insights into the vemurafenib resistance mechanisms in A375 cells, we performed Gene Ontology (GO) enrichment analysis with the top 33 genes that regulate resistance to vemurafenib using the online tool DAVID [19], [20]. Many GO terms under Biological Process were redundant under different annotation clusters (Table S6). We removed the redundant GO terms using the online tool REVIGO [21]. Visualization of GO data by REVIGO (Figure 4B) indicated that genes that modulate vemurafenib resistance are involved in histone/protein acetylation, chromosome/chromatin organization, cell cycle, protein complex biogenesis, DNA template transcription/initiation and regulation of transcription from RNA Pol II promoter. Although validation of top hits with targeted knockouts is necessary, we discuss below how the identified hit genes might regulate resistance to vemurafenib on the basis of information available in the literature.

### MAPK pathway

Four genes NF1, MED12, CUL3 and NF2 involved in MAPK signalling pathway were among our top 11 hits. These genes were obtained in previous screens in agreement with the notion that activation of MAPK signalling pathway is the main mechanism leading to vemurafenib resistance.

NF1 (Neurofibromin 1), a large protein consisting of over 2800 amino acid residues, contains a domain similar to the catalytic domain of GTPase Activating Protein (GAP). NF1 negatively regulates RAS by stimulating its GTPase activity [22]. Therefore, loss of NF1 would result in activation of Ras and MAPK signalling pathways.

CUL3 is a subunit of multiple E3 ubiquitin ligases and its loss is expected to cause stabilization of several proteins. CUL3 was recently shown to enhance MAPK signalling by stabilization and Src-dependent activation of RAC1 [23]. Interestingly, CUL3 is also required for proteasomal degradation of NF1 [24] but its effect on RAC1 appears to override its effect of NF1 stability in terms of MAPK signalling and vemurafenib resistance [23].

NF2 encodes merlin which is mutated in various forms of cancer such as schwannomas, mesotheliomas, breast, prostate, colorectal, hepatic, clear cell renal cell carcinoma and melanomas [25]. Loss of merlin was reported to cause vemurafenib resistance and the presence of merlin even at very low levels was sufficient to maintain vemurafenib sensitivity [25]. Merlin inhibits Rac by preventing its localization to the plasma membrane [26] and by inhibiting the p21-activated kinase PAK1 [27]. So inactivation of merlin would result in Rac activation and stimulation of the MAPK pathway. Loss of merlin activates other growth promoting pathways such as Hippo and mTOR [25] explaining the incidence of NF2 mutations in several forms of cancers.

MED12 is a part of the Transcriptional MEDIATOR complex that physically binds to TGF-βR2 and inhibits the TGF-βR2 signalling pathway. MED12 suppression therefore results in activation of TGF-βR signalling leading to increased RAS-MEK-ERK signalling. Mutations in MED12 confer resistance to a number of drugs used against colon cancer, melanoma and liver cancer [28]. We also identified 4 additional subunits of the MEDIATOR complex, namely MED10, MED15, MED19 and MED23 which have all been reported in previous genetic screens [12][11][13].

### Epigenetic regulation

Post-translational histone modifications such as acetylation, methylation and sumoylation have significant effects on gene expression. Histone acetylation is commonly associated with activation of gene expression whereas histone methylation is linked to either activation or repression of gene expression [28]. Studies of epigenomic alterations revealed a loss of histone acetylation and histone H3 Lys 4 methylation (H3K4me2/3) on regulatory regions proximal to specific cancer-regulatory genes involved in important signalling pathways driving melanoma [14]. We identified several genes related to post-translational histone modification in our screen namely TAF6L, TAF5L, CCDC101, SUPT20H, TADA2B, TADA1, TADA3, HIRA and SMARCA4.

Seven genes (TAF6L, TAF5L, CCDC101, SUPT20H, TADA2B, TADA1 and TADA3) encode subunits of the SPT3-TAFII31-GCN5L acetylase (STAGA complex). STAGA is a chromatin-acetylating transcription coactivator and regulates numerous cellular processes like transcription, splicing and DNA repair through coordination of multiple histone post-translational modifications [29], [30].

Apart from the STAGA complex, factors mediating chromatin organization such as histone chaperones (HIRA) and SMARCA4 (SWI/SNF related, matrix associated, actin dependent regulator of chromatin, subfamily a, member 4) were among our top hits. Mutations in SWI/SNF chromatin remodelling genes have been associated with invasive melanomas [31]. SMARCA4 promotes chromatin accessibility around double-strand breaks (DSBs) [32] and HIRA is recruited to DSBs, facilitating restoration of chromatin structure by depositing histones [33]. Therefore, loss of HIRA and SMARCA4 is expected to cause genomic instability, a hallmark of tumour cells [34].

### Cell cycle

Dysregulation of cell cycle can boost cell division rates by overriding cell cycle arrests, inhibiting apoptosis and by promoting genetic instability. We found four genes related to cell cycle in our screen namely FOXD3, ANAPC11, PELO and MAU2 with the last three being novel hits.

FOXD3 (Forkhead transcription factor) was previously reported to play a role in resistance towards the precursor of vemurafenib, PLX4720 [35]. As a potent antagonist of melanoma proliferation [36], FOXD3 prevents melanoma cell migration and invasion in a Rho-associated protein kinase dependent manner [37]. Mutant BRAF signalling results in FOXD3 downregulation [36], which was also linked to epithelial–mesenchymal transition (EMT)-like phenotype [38]. Attenuation of mutant BRAF signalling caused increased FOXD3 levels [36]. Overexpression of FOXD3 inhibited growth of melanoma cells by causing a p53-dependent cell cycle arrest [36]. Deletion of FOXD3 in vemurafenib-treated A375 cells could therefore overcome the cell cycle arrest and result in vemurafenib-resistance.

MAU2 interacts with NIPBL/SCC2 to form a heterodimeric complex that is required for loading of cohesin complex onto chromatin [39], [40]. This enables cohesin to tether sister chromatids together immediately after their generation during DNA replication until their separation during anaphase. ANAPC11 is the catalytic subunit of the anaphase promoting complex/cyclosome (APC/C) that regulates progression through mitosis and the G1 phase of the cell cycle. Loss of MAU2 and ANAPC11 could enhance genetic instability which could accelerate the acquisition of mutations causing vemurafenib resistance.

### Translation

Misregulation of protein translation is a key mechanism that drives cellular transformation and tumour growth [41]. We found that 5 of the top 33 hit genes encode proteins involved in translation, with 3 of them being mitochondrial. The 5 genes encode PELO (ribosome rescue/mRNA surveillance factor), POLR3K (DNA-dependent RNA polymerase involved in 5S rRNA and tRNA synthesis), GFM1 (G elongation Factor Mitochondrial 1), MRPS12 (Mitochondrial Ribosomal Protein S12) and MARS2 (mitochondrial Methionine-tRNA synthetase 2).

PELO is an evolutionarily conserved gene [42] required for ribosomal disassembly [43]. PELO is involved in the no-go decay (NGD) surveillance mechanism which leads to mRNA degradation in stalled ribosomal elongation complexes (ECs) [44]. PELO knockdown has been reported to activate PI3K/AKT [45], [46] and BMP (Bone Morphogenetic Protein) signalling pathways [46], which resulted in epidermal hyperplasia in mice [46]. Furthermore, conditional deletion of PELO in mouse epidermal stem cells resulted in their hyperproliferation [47].

Notably, three mitochondrial translation proteins (GFM1, MRPS12 and MARS2) were among the top 33 hits. BRAF mutant cells have increased rates of glycolysis and reduced oxidative phosphorylation in comparison to wild type cells [48]. Treatment of BRAF mutant cells with vemurafenib reduces glycolytic flux and increases mitochondrial oxidative phosphorylation resulting in severe oxidative stress [49]. It would be informative to test whether inhibition of mitochondrial translation relieves oxidative stress in vemurafenib-treated BRAF mutant cells thereby providing a fitness advantage.

### Other cellular processes

We obtained AHR (Aryl Hydrocarbon/dioxin receptor) which encodes a ligand-dependent transcription factor in our screen. In response to agonist binding, AHR translocates to the nucleus and activates the expression of xenobiotic metabolism genes like the P450 family of enzymes. Intriguingly, stable activation of AHR was reported to be required for resistance to BRAF-inhibitors in melanoma [50]. Vemurafenib was shown to bind to a non-canonical substrate binding site of AHR and inhibit the canonical AHR signalling pathway [50]. Our result appears to be at odds with this observation. However, AHR was shown to have either oncogenic or tumour suppressive effects depending on cellular phenotype [51]. AHR knockdown promoted melanoma in mouse [51]. In addition, the AHR levels in human metastatic melanomas were reduced in comparison to benign nevi [51].

Another novel hit Ctc1 is a component of the CST (Ctc1, Stn1 and Ten1) complex, which functions as a terminator of telomerase activity [52]. Depletion of Cst1 boosted telomerase activity and telomere elongation [52]. Interestingly, downregulation of the Ras pathway by inhibition of MEK/ERK kinases promoted telomere DNA damage and fragility in aggressive lung cancer and glioblastoma (GBM) mouse models [53]. It is conceivable that increased telomere length caused by deletion of Ctc1 might provide a growth advantage in vemurafenib-treated cells.

Our list of hits included CYC1 (mitochondrial ubiquinol-cytochrome c oxidoreductase: functions in the Electron Transport Chain), CYP2R1 (Cytochrome P450 family 2 subfamily R member 1: a 25-hydroxylase involved in Vitamin D3 metabolism), CAND1 (cullin-associated neddylation-dissociated protein 1: a F-Box exchange factor), VBP1 (VHL binding protein 1: binds to E3 Ubiquitin ligase Von Hippel Lindau), PGD1 (phosphogluconate dehydrogenase: Enzyme in the pentose phosphate pathway) and C7orf26 (Chromosome 7 Open Reading Frame 26: Uncharacterized protein). All of these hits except CAND1 and PGD1 are novel.

In summary, genome-scale screen for modulators of vemurafenib resistance in A375 cells with the Brunello library identified 19 previously reported genes and 14 novel hits. Confirming and dissecting how inactivation of novel genes results in vemurafenib resistance might generate new therapeutic strategies for countering melanoma. Newly identified hits can potentially serve as melanocytic biomarkers for cancer detection.

## Acknowledgements

This research was supported by the Agency for Science, Technology and Research (A*STAR) under its Industry Alignment Fund – Pre-Positioning Programme (IAF-PP) grant number H18/01/a0/B14 as part of the A*STAR Innovations in Food and Chemical Safety (IFCS) Programme.

## Author Contributions

CJHG and JHW performed the CRISPR-based genetic screen. CEF, CJHG and JHW did the analysis of the NGS data. BXT and CC supervised the lentiviral transfection experiments. YHL and DL supervised the bioinformatics and CRISPR experiments respectively. PA, JHW and CJHG wrote the MS. PA conceived and supervised the entire project.

## Supplementary information

**Figure S1:**
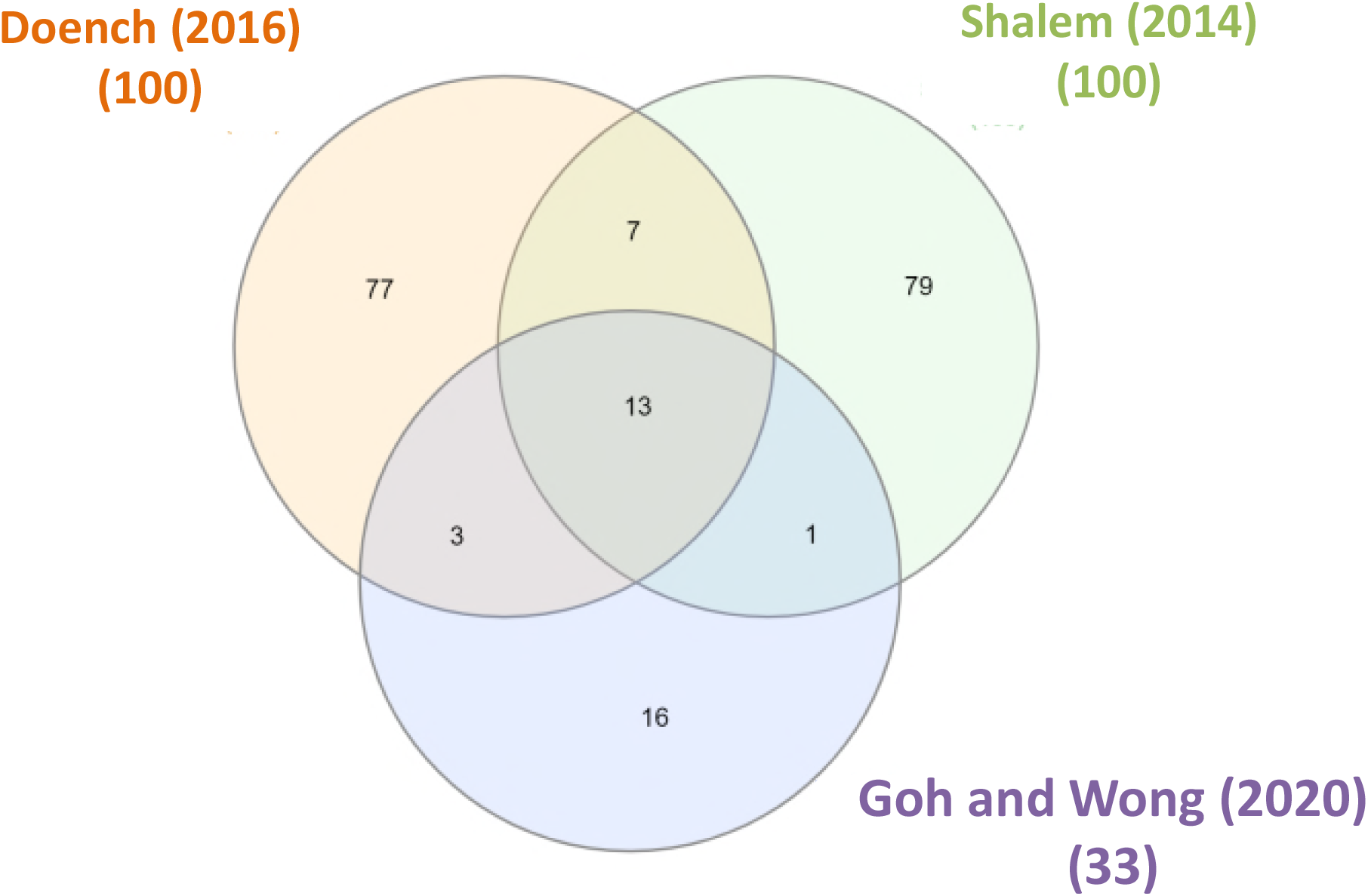
Venn diagram of top 33 hits versus top 100 hits from Shalem *et al.* (2014) and Doench *et al.* (2016).

**Table S1: Primer sequences for amplifying sgRNA Brunello library and NGS**.

The NGS-Lib-Fwd primers contain 1-10bp staggered nucleotides and the NGS-Lib-Rev primers provide unique barcodes. Bold letters are the unique barcodes of the reverse primers.

**Table S2: Read count file of the 76,441 sgRNA in the Brunello library targeting of the 19,114 genes**.

**Table S3: MAGeCK Enriched genes rank after 14 days of vemurafenib treatment**.

This excel file contains two tabs, where the first tab contains all the MAGeCK Enriched genes rank of 19,114 genes and the second tab only shows the MAGeCK Enriched genes rank with the p-value less than 0.05 and the read count of the all 4 sgRNAs for each gene are shown.

**Table S4: Comparison of the top 33 hits with top 100 enriched genes in Whittaker *et al.* (2013), Wei Li *et al.* (2014), Doench *et al.* (2016), Sage *et al.* (2017) and Sanson *et al.* (2018)**.

**Table S5: STRING analysis of the top 33 genes that regulate vemurafenib resistance**.

**Table S6: Gene Ontology (GO) analysis of the top 33 genes that regulate vemurafenib resistance using the online tool DAVID and REVIGO visualization**.

## Notes

### Competing Interest Statement

The authors have declared no competing interest.

