## Supplementary tables for "Identification of pathways modulating Vemurafenib resistance in melanoma cells via a genome-wide CRISPR/Cas9 screen": Table S1_Primer list.docx

**Table S6: Primer sequences for amplifying sgRNA Brunello library and NGS.**

| **Primer names** | **5' -> 3'** |
| --- | --- |
| NGS-Lib-Fwd-1 | AATGATACGGCGACCACCGAGATCTACACTCTTTCCCTACACGACGCTCTTCCGATCTTAAGTAGAGGCTTTATATATCTTGTGGAAAGGACGAAACACC |
| NGS-Lib-Fwd-2 | AATGATACGGCGACCACCGAGATCTACACTCTTTCCCTACACGACGCTCTTCCGATCTATCATGCTTAGCTTTATATATCTTGTGGAAAGGACGAAACACC |
| NGS-Lib-Fwd-3 | AATGATACGGCGACCACCGAGATCTACACTCTTTCCCTACACGACGCTCTTCCGATCTGATGCACATCTGCTTTATATATCTTGTGGAAAGGACGAAACACC |
| NGS-Lib-Fwd-4 | AATGATACGGCGACCACCGAGATCTACACTCTTTCCCTACACGACGCTCTTCCGATCTCGATTGCTCGACGCTTTATATATCTTGTGGAAAGGACGAAACACC |
| NGS-Lib-Fwd-5 | AATGATACGGCGACCACCGAGATCTACACTCTTTCCCTACACGACGCTCTTCCGATCTTCGATAGCAATTCGCTTTATATATCTTGTGGAAAGGACGAAACACC |
| NGS-Lib-Fwd-6 | AATGATACGGCGACCACCGAGATCTACACTCTTTCCCTACACGACGCTCTTCCGATCTATCGATAGTTGCTTGCTTTATATATCTTGTGGAAAGGACGAAACACC |
| NGS-Lib-Fwd-7 | AATGATACGGCGACCACCGAGATCTACACTCTTTCCCTACACGACGCTCTTCCGATCTGATCGATCCAGTTAGGCTTTATATATCTTGTGGAAAGGACGAAACACC |
| NGS-Lib-Fwd-8 | AATGATACGGCGACCACCGAGATCTACACTCTTTCCCTACACGACGCTCTTCCGATCTCGATCGATTTGAGCCTGCTTTATATATCTTGTGGAAAGGACGAAACACC |
| NGS-Lib-Fwd-9 | AATGATACGGCGACCACCGAGATCTACACTCTTTCCCTACACGACGCTCTTCCGATCTACGATCGATACACGATCGCTTTATATATCTTGTGGAAAGGACGAAACACC |
| NGS-Lib-Fwd-10 | AATGATACGGCGACCACCGAGATCTACACTCTTTCCCTACACGACGCTCTTCCGATCTTACGATCGATGGTCCAGAGCTTTATATATCTTGTGGAAAGGACGAAACACC |
| NGS-Lib-KO-Rev-1 | CAAGCAGAAGACGGCATACGAGAT**TCGCCTTG**GTGACTGGAGTTCAGACGTGTGCTCTTCCGATCTCCGACTCGGTGCCACTTTTTCAA |
| NGS-Lib-KO-Rev-2 | CAAGCAGAAGACGGCATACGAGAT**ATAGCGTC**GTGACTGGAGTTCAGACGTGTGCTCTTCCGATCTCCGACTCGGTGCCACTTTTTCAA |
| NGS-Lib-KO-Rev-3 | CAAGCAGAAGACGGCATACGAGAT**GAAGAAGT**GTGACTGGAGTTCAGACGTGTGCTCTTCCGATCTCCGACTCGGTGCCACTTTTTCAA |
| NGS-Lib-KO-Rev-4 | CAAGCAGAAGACGGCATACGAGAT**ATTCTAGG**GTGACTGGAGTTCAGACGTGTGCTCTTCCGATCTCCGACTCGGTGCCACTTTTTCAA |
| NGS-Lib-KO-Rev-5 | CAAGCAGAAGACGGCATACGAGAT**CGTTACCA**GTGACTGGAGTTCAGACGTGTGCTCTTCCGATCTCCGACTCGGTGCCACTTTTTCAA |
| NGS-Lib-KO-Rev-6 | CAAGCAGAAGACGGCATACGAGAT**GTCTGATG**GTGACTGGAGTTCAGACGTGTGCTCTTCCGATCTCCGACTCGGTGCCACTTTTTCAA |
| NGS-Lib-KO-Rev-7 | CAAGCAGAAGACGGCATACGAGAT**TTACGCAC**GTGACTGGAGTTCAGACGTGTGCTCTTCCGATCTCCGACTCGGTGCCACTTTTTCAA |
| NGS-Lib-KO-Rev-8 | CAAGCAGAAGACGGCATACGAGAT**TTGAATAG**GTGACTGGAGTTCAGACGTGTGCTCTTCCGATCTCCGACTCGGTGCCACTTTTTCAA |
| NGS-Lib-KO-Rev-9 | CAAGCAGAAGACGGCATACGAGAT**AAGTAGAG**GTGACTGGAGTTCAGACGTGTGCTCTTCCGATCTCCGACTCGGTGCCACTTTTTCAA |
